# Synaptic mechanisms of context-dependent sensory responses in the hippocampus

**DOI:** 10.1101/624262

**Authors:** Xinyu Zhao, Yingxue Wang, Nelson Spruston, Jeffrey C. Magee

**Affiliations:** Howard Hughes Medical Institute, Janelia Research Campus, Ashburn, VA, 20147; Max Planck Florida Institute for Neuroscience, 1 Max Planck Way, Jupiter, FL 33458; Howard Hughes Medical Institute, Department of Neuroscience, Baylor College of Medicine, Houston, TX, 77003

## Abstract

As animals navigate, they must identify features in context. The hippocampus-a structure critical for navigation-exhibits a different code for each environment, uniquely representing locations and objects within each environment in the firing rates of its neurons. It is unknown how or where these context-specific codes emerge. For example, it is unknown whether pyramidal neurons in the CA1 subregion produce these codes (by combining separate inputs from local sensory cues and the environmental context) or inherit conjunctive codes from upstream brain areas. We performed electrical recordings in the hippocampus as mice navigated in two distinct virtual environments. In CA1, subthreshold synaptic responses to a visual cue in one environment were absent when the same cue occurred in the second environment. In CA3, cue-driven spiking also strongly depended on environmental context. These results indicate that context-dependent sensory coding in CA1 is inherited from its CA3 input.

## Introduction

Position-variant sensory cues constitute a set of spatial landmarks that guide animals during spatial navigation. To do so, sensory input is processed at multiple locations in the brain. Responses in early sensory systems are predominantly evoked by relatively simple features of instantaneous stimuli within localized receptive fields, while higher sensory areas encoded more complex features. At even higher levels (e.g. hippocampus), it is ethologically beneficial for a navigation processor to modulate sensory responses to a given cue depending on the particular context since such encoding can prevent confusion when similar objects occur in distinct environments. It is thus important to understand to what extent the brain can differentiate the same sensory cue in different contexts and how such a computation is implemented.

Behavioral and physiological studies have identified the hippocampus as a critical brain area encoding space^1^. Pyramidal cells in the hippocampus have been found to selectively fire action potentials when an animal enters a specific location within the environment^2^ and it has been shown that sensory cues exhibit substantial influence on the firing of these neurons. For example, in rodents, the firing fields of places cells follow the rotation of visual landmarks^3, 4^, and in primates, ‘spatial view cells’ in the hippocampus become active when the animal’s eyes are fixing at a particular object, indicating a strong visual sensory drive^5–7^.

Several lines of evidence have suggested that sensory responses in the hippocampus might be modulated by environmental context. Behavioral studies have shown that the hippocampus is indispensable in contextual fear conditioning in which animals were trained to recognize a specific context where they received an aversive stimulus. However, hippocampus is not required in tasks where the aversive stimulus was only associated with a single cue^8^, implying that the hippocampus might play an essential role in encoding the global context rather than individual cues. Consistently, physiological recordings have revealed that in some hippocampal neurons, responses produced by local objects depend on the object’s position relative to other cues^9, 10^. In addition, place-field locations are largely remapped when animals are placed in differently shaped chambers containing the same salient cue card^11, 12^. Although these experiments are informative, they face the potential confound that the level of context dependence exhibited may depend on how isolated the sensory cues were from one another. Indeed, multiple cues, across different modalities, can usually be perceived together by animals at a given location making it difficult to completely alter all sensory features associated with one location. In addition, some cues, especially visual ones, can be accessible to the animal across a wide range of locations. Therefore, the manipulation of a local object may not only change the sensory input when the animal gets close to it, but also the input at many other locations. Thus, direct assessment of the context-dependence of place-cell responses requires full control and separation of sensory cues in space, so that one can independently manipulate instantaneous and preceding sensory cues.

In addition, it is still unclear from previous studies where in the hippocampal circuit the input from one sensory cue is integrated with context. Most studies of context-dependent place-cell firing have focused on the CA1 subregion, which receives multiple excitatory inputs, including inputs from CA3, CA2, entorhinal cortex (EC), thalamus, and septal nuclei. One possibility is that signals evoked by a cue and its context may be channeled into CA1 through separate pathways^13–17^. Alternatively, contextual and sensory inputs may be integrated in upstream areas (e.g. the CA3 subregion) with the conjunctive tuning largely inherited by CA1. Previous extracellular recordings, which can only measure firing rates, do not provide information about how synaptic input to a single cell is altered following manipulation of the environmental context.

To address these questions, we combined a virtual reality system^18^, which facilitates strict control and flexible real-time manipulation of visual cues, with in-vivo intracellular recordings, thus permitting the measurement of subthreshold membrane potential as well as spiking. We found that the firing fields of CA1 place cells closely followed the movement of a visual cue, regardless of its position within one environment. Strikingly, however, the same cue induced dramatically different responses when presented in a distinct environment. Membrane potential dynamics suggested that context-dependent responses in CA1 resulted from remapping of its excitatory inputs. Consistently, extracellular recordings in the CA3 subregion revealed that CA1 receives context-dependent sensory information from its CA3 input.

## Results

### Induction of place fields by behavioral timescale synaptic plasticity in CA1 pyramidal cells during virtual navigation

We trained head-fixed male mice to run on a spherical treadmill, modified from a previous design^19^. The visual scene was rendered on three monitors around the animal, covering 216° of its visual field, and movement of the virtual world was coupled to locomotion through a customized virtual reality system (Fig. 1a). Mice were trained to run clockwise on a 150cm virtual oval track comprised of two straight arms and two 180° turns (Fig. 1b). The track was decorated with six distinguishable visual patterns at different locations (Fig. 1b and Supplementary Fig. 1a). Visual cues at two locations—vertical and horizontal gratings on opposite sides of the track (cues A and D, locations 1 and 4, Fig. 1b)—were manipulated throughout this study. Since we aimed to investigate sensory integration in the hippocampus, non-sensory factors known to affect the hippocampal activity, including reward anticipation^20^ and locomotion speed^21, 22^, were controlled by two symmetrical reward zones on the track (Fig. 1b,c). With this reward paradigm, running speed was similar around the two manipulated locations (Supplementary Fig. 2a). Although animals were not required to take any actions to trigger the water drop in the reward zones, they typically slowed and performed anticipatory licking prior to reward delivery, indicating that the association between the position and reward was learned (Fig. 1c, d; Supplementary Fig. 2a).

**Figure 1.**
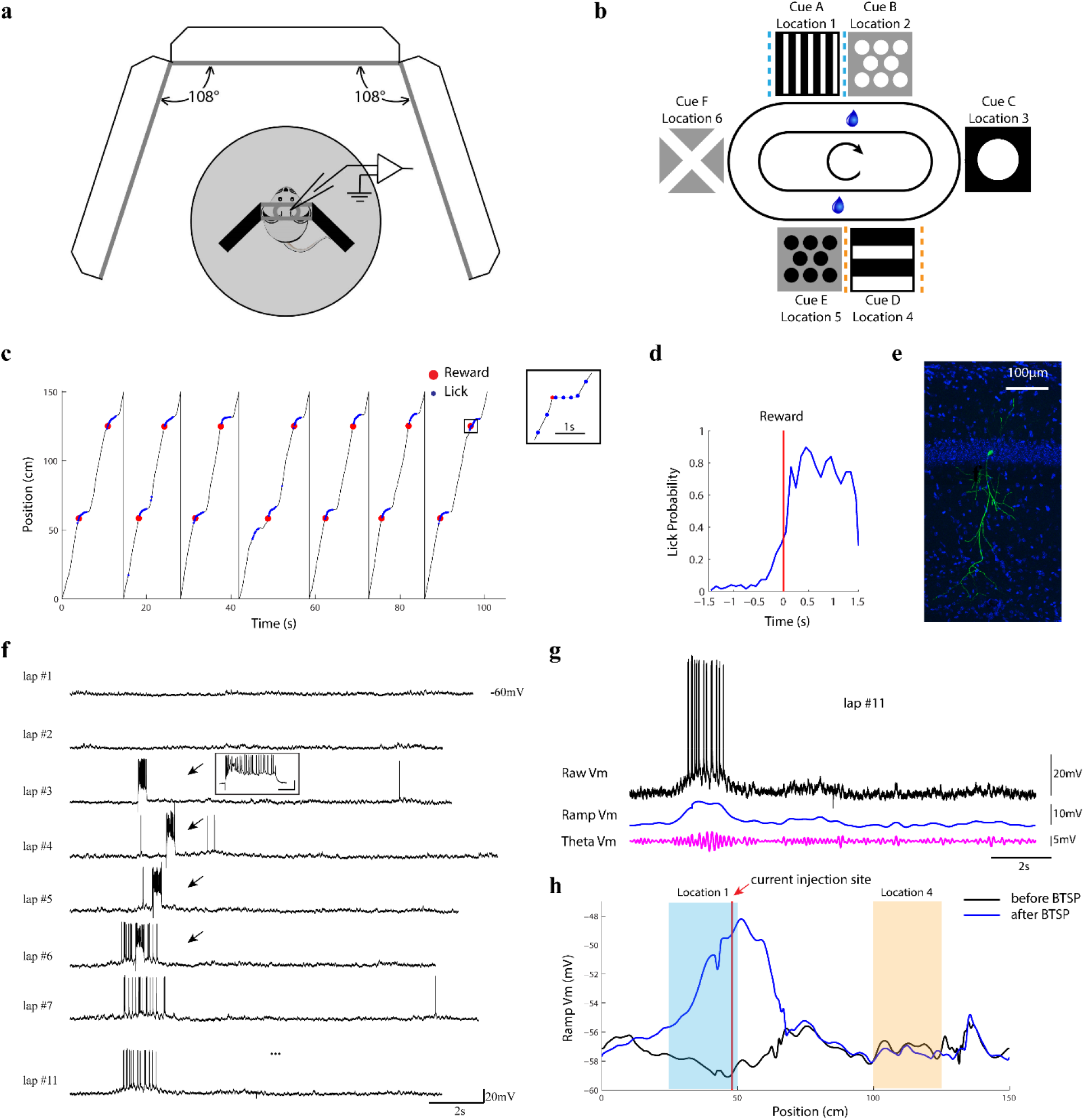
Induction of place fields by BTSP during virtual navigation. **(a)** Mice running on a spherical treadmill to navigate in a virtual reality environment. **(b)** Configuration of the virtual oval track. Six visual cues were presented at different locations along the track. Two reward zones were located at the center of each straight arm. Two cues that were manipulated throughout this study, A (vertical grating) and D (horizontal grating), are marked by light blue and yellow dashed lines, respectively. **(c)** Behavior of an example animal. Inset: zoom-in view around one reward delivery to show that licking (blue) started before the reward (red). **(d)** Anticipatory licking in the example animal as in (c). Averaged lick probability was plotted against the time relative to each reward delivery. **(e)** An example CA1 pyramidal cell (green) labeled by the whole-cell recording (blue: DAPI). **(f)** BTSP induced a stable place field in an originally silent CA1 cell. Arrows indicate square step current injections. Inset: plateau potentials induced by the current injection (scale bars: 100ms, 30mV). **(g)** Subthreshold ramp Vm (blue) and theta oscillation (magenta) were extracted by band-pass filters (<2Hz for ramp; 4-10Hz for theta) from the recorded membrane potential. Note that the ramp depolarization is accompanied by an increase in theta amplitude. **(h)** Position-variant ramp depolarization before (black) and after (blue) BTSP. Current injection position during the BTSP induction is labeled by the red line. Light blue and yellow shadings mark the locations of cue A and D, respectively, as in (b).

To assess not only the output but also the input of hippocampal place cells, we recorded membrane potential (V_m_) in CA1 pyramidal cells with whole-cell patch-clamp recordings (Fig. 1e). Previous studies have shown that most CA1 cells are not active during virtual navigation^19, 23, 24^; and those that are active may exhibit place fields with diverse sizes and locations. Therefore, with blind patching, the chance of recording from a place cell that can respond to one cue in a specific virtual environment is very low. Fortunately, we recently discovered that new place-cell responses can be generated by synaptic plasticity. Specifically, dendritic plateau potentials create new place fields by potentiating position-tuned synaptic inputs via a mechanism called “behavioral timescale plasticity (BTSP) ^25, 26^. BTSP occurs naturally through spontaneous plateau potentials, but can also be induced experimentally. We therefore induced place fields by injecting current to evoke plateau potentials at the locations of the manipulated cues^26^. In the first set of experiments, twelve CA1 cells, initially without firing fields during the task, were recorded. Step current injections (300 ms, 600-800 pA) were triggered at the end of location 1 (cue A) for 4-10 consecutive laps to generate plateau potentials and related synaptic potentiation (Fig. 1f). Consistent with our previous findings in mice running on a linear belt, the induction protocol created a place field near the current injection site, which persisted throughout the entire recording session (Fig. 1f-1h). The induced field recapitulated all three major aspects of intracellularly recorded natural place fields, including a slow ramp depolarization, a larger theta oscillation and a significant spiking field (Fig. 1g). Cue A was located in the early half of the resulting field, covering the rising phase of the V_m_ depolarization ramp (see Fig. 1h as an example and 2c as the population average).

### Place fields tightly followed sensory cue manipulations

To investigate the impact of sensory cues on place fields, after a stable field was established in a recorded cell, we duplicated cue A on the opposite side of the track (i.e., cue D was replaced by cue A at location 4; Fig 1b). Following cue duplication, CA1 place cells fired at both instantiations of cue A; however, the response at the new location was smaller. All three metrics (ramp depolarization, theta amplitude and spiking rate) indicated that the activity at location 4 was significantly enhanced compared to the control condition, but significantly weaker than the original field at location 1 (Fig 2c-2h; ramp: 10.5±1.5mV vs. 6.3±1.3mV; theta: 2.8±0.3mV vs. 1.9±0.3mV; rate: 16.8±1.8 Hz vs. 9.3±3.2 Hz; original vs. new fields). On the other hand, the activity at the original location of cue A (location 1) was not significantly affected (Fig 2c-2h; ramp: 10.5±0.8 mV vs. 10.5±1.5 mV; theta: 2.9±0.2 mV vs. 2.8±0.3 mV; rate: 16.8±1.8Hz vs. 20.2±3.2Hz; before vs. after cue duplication).

**Figure 2.**
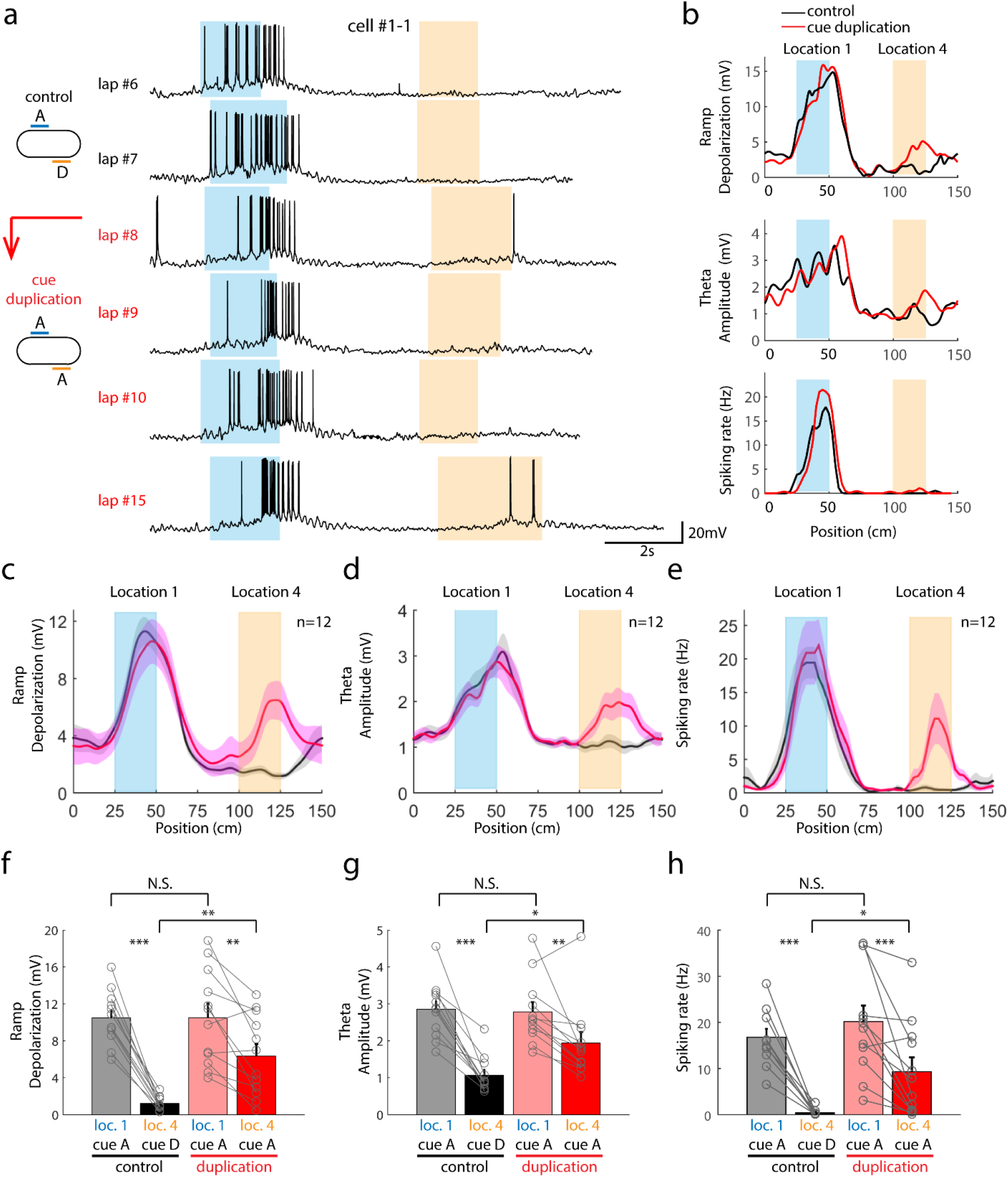
Cue duplication induced a second place field with smaller amplitude. **(a)** An example cell with the duplication of cue A during recording. **(b)** Ramp depolarization, theta amplitude and spiking rate before (black) and after (red) the cue duplication in the example cell shown in(a). **(c)-(e)** Position-variant ramp depolarization (c), theta amplitude (d) and spiking rate (e) before (black) and after (red) the cue duplication (n=12 cells from 7 animals). Solid curves and shaded areas depict mean±s.e.m. Through (b) to (e), locations of cue A and D were marked with light blue and yellow shadings, respectively, as in Fig. 1. **(f)-(h)** Statistical comparisons of peak ramp depolarization (f), theta amplitude (g) and spiking rate (h) following the cue duplication. Peak values of each metric were calculated within location 1 (between the two light blue lines in (c)-(e); grey for control, pink for cue duplication) and location 4 (between the two yellow lines in (c)-(e); black for control, red for cue duplication). Ramp depolarization (f): P=1.485e-7 for control loc. 1 vs. control loc. 4, 0.0027 for duplication original vs. duplication new, 0.9783 for control loc. 1 vs. duplication loc. 1, and 0.001 for control loc. 4 vs. duplication loc. 4. Theta amplitude (g): P=1.029e-5 for control loc. 1 vs. control loc. 4, 0.0061 for duplication loc. 1 vs. duplication loc. 4, 0.7968 for control loc. 1 vs. duplication loc. 1, and 0.0116 for control loc. 4 vs. duplication loc. 4. Spiking rate (h): P=8.163e-7 for control loc. 1 vs. control loc. 4, 0.0024 for duplication loc. 4 vs. duplication loc. 4, 0.1988 for control loc. 1 vs. duplication loc. 1, and 0.0083 for control loc. 4 vs. duplication loc. 4. Paired student t-test was conducted in all analyses. *: P<0.05, **: P<0.01, ***: P<0.001, N.S.: not significant (P>=0.05).

To determine if the partial duplication of place fields in this experiment arises from incomplete manipulation of visual input (i.e. animals seeing adjacent unmanipulated cues), we added two ‘virtual curtains’, which animals could run through but not see through, at the center of each straight arm (Supplementary Fig. 1b). As a result, the animal’s visual inputs were limited to cue A or B at their respective locations. Virtual curtains were present from the beginning of training and did not disrupt the animal’s locomotion (Supplementary Fig 2a). With visual barriers in place, duplication of cue A produced a full-sized place field at the new location (Ramp: 11.7±1.4 mV vs. 11.2±1.2 mV; Theta; 3.0±0.3 mV vs 2.9±0.2 mV; rate: 17.3±3.7 vs 16.2±3.0 Hz; location 1 vs. location 4 after cue duplication; Fig. 3a-c; Supplementary Fig 3). These results demonstrate that place-specific sensory cues are sufficient to drive place-cell firing.

**Figure 3.**
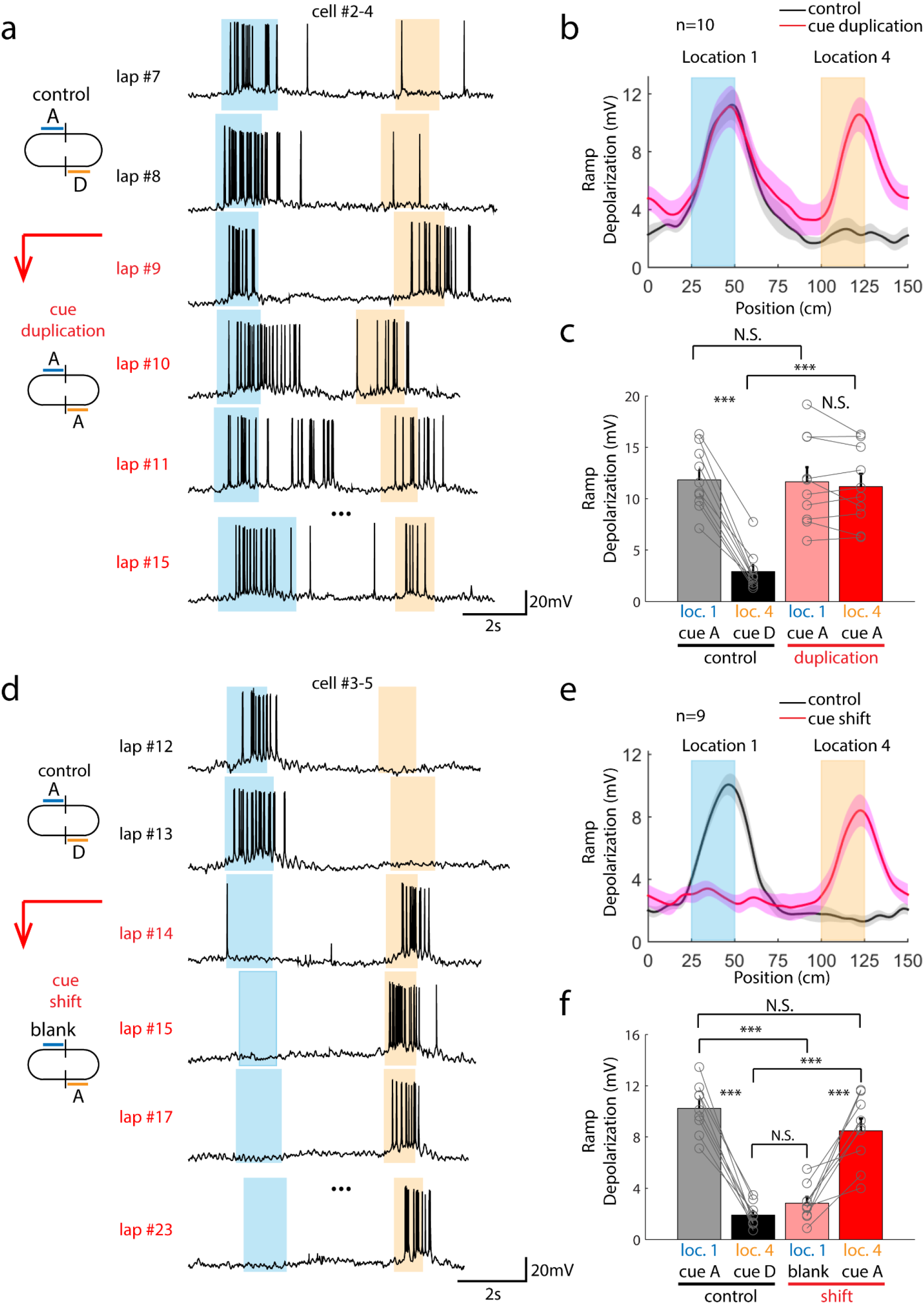
Cue isolation by visual barriers revealed strong control of visual cues on CA1 place fields. **(a)** An example cell with the duplication of isolated cue A during recording. **(b)** Position-variant ramp depolarization before (black) and after (red) the cue duplication (n=10 cells from 7 animals). **(c)** Statistics of peak ramp depolarization shown in (b): P=8.744e-7 for control loc. 1 vs. control loc. 4, 0.3156 for duplication loc. 1 vs. duplication loc. 4, 0.8822 for control loc. 1 vs. duplication loc. 1, and 1.159e-5 for control loc. 4 vs. duplication loc. 4. **(d)** An example cell with the shift of isolated cue A during recording. **(e)** Position-variant mean ramp depolarization before (black) and after (red) the cue shift (n=9 cells from 7 animals). **(f)** Statistics of peak ramp depolarization shown in (e): P=2.12e-6 for control loc. 1 vs. control loc. 4, 8.333e-5 for shift loc. 1 vs. shift loc. 4, 1.726e-5 for control loc. 1 vs. shift loc. 1, 0.051 for control loc. 1 vs. shift loc. 4, 4.258e-4 for control loc. 4 vs. shift loc. 4, and 0.1427 for control loc. 4 vs. shift loc. 1. Paired student t-test was conducted in all analyses. All color conventions and statistics labels are the same as in Fig. 2.

We next sought to test whether the place-specific sensory cue is also necessary for place-field firing. In a second set of recordings, cue A was moved from its original location to the opposite side of the track (i.e., cue A moved from location 1 to location 4, with only black walls displayed at location 1 after the shift). All three metrics of place cells showed that place fields had moved to the new location, immediately and at full amplitude, following this cue shift (Ramp: 10.2±0.8 mV vs. 8.5±1.0 mV; Theta; 3.4±0.2 mV vs. 2.9±0.3 mV; rate 19.7±3.2 Hz vs. 16.3±3.9 Hz; control location 1 vs. location 4, Fig. 3d, e; Supplementary Fig. 3). However, the activity at the original location was reduced essentially to the baseline (Fig. 3e: ramp: 1.9±0.3 mV vs. 2.8±0.5 mV; theta: 3.2±0.2 mV vs 2.8±0. 2mV; rate: 22.7±4.8 Hz vs. 20.2±5.3 Hz; control location 1 vs. location 4, p>0.05; Supplementary Fig. 3), indicating that the visual cue fully accounts for the field. It should be noted that the duplicated/shifted place fields fully emerged on the first lap immediately after cue manipulations, indicating that the observed effects reflect direct responses without a requirement for additional plasticity (Figure 3a and 3d as examples).

In the above experiments two reward zones were located adjacent to both cues A and D, thus making these two cues potentially special to the animal. To control for any cue-reward association effects we recorded CA1 cells during a random reward-delivery paradigm. As expected, animals constantly licked to check the reward availability, and exhibited approximately uniform running speed along the track (Supplementary Fig. 2b and Supplementary Fig. 4a). Cue duplication under the random-reward condition showed identical effects as under the fixed reward condition (Supplementary Fig. 4b-4g), indicating that the reward association does not influence the results.

CA1 receives synaptic inputs from multiple brain areas. Which pathway primarily determines the sensory representation in CA1 in our BTSP-induced place fields? Although anatomical and in-vitro physiological studies have suggested that CA3 dominates the excitatory input to CA1^27–30^, several in-vivo studies have challenged the idea that CA3 can provide the sensory information^17, 31^. Instead, it has been proposed that the sensory input may be conveyed by EC or CA2. To test whether CA1 receives the relevant sensory information from CA3, we performed extracellular recordings in CA3 using 64-channel silicon probes in chronically implanted adult mice that underwent the same virtual reality behavioral training described above (see Methods for details). Only CA3 place cells with stable place fields during recording sessions were included in the analysis (Supplementary Fig. 5, see Methods for details).

Movement of visual cues impacted the firing of CA3 place cells around the locations where cues were changed (see example recording sessions in Supplementary Fig. 5e, f). To examine this effect, we first identified place fields from rate maps (Supplementary Fig. 6), and then selectively analyzed those place fields whose firing overlapped the manipulated cue for at least half of the cue length (overlapped length>12.5cm). As seen in individual cells (Fig. 4a) and the population average (Fig. 4b), cue duplication produced a second field with little difference to the original field. The average spiking rate at the manipulated location was significantly increased after addition of the duplicated cue (averaged spiking rate: 4.1±1.1 Hz vs. 0.3±0.1 Hz, control field 2 vs. duplication field 2 in Fig. 4c), and was not significantly different from the firing rate at the location of the original place field (averaged spiking rate: 4.5±1.0 Hz vs. 4.1±1.1 Hz, control field 1 vs. duplication field 1 in Fig. 4c). The cue shift reduced place field activity at the cue’s original location, although the extent of the reduction varied across cells (Fig. 4d). On the population level, CA3 activity at the cue location was significantly reduced by the cue shift (Fig. 4e, f; averaged spiking rate: 5.0±1.0 Hz vs. 1.7±0.3 Hz, control field 1 vs. shift field 1), although the remaining activity was slightly higher than the baseline (Fig. 4e, f; averaged spiking rate: 0.6±0.2 Hz vs. 1.7±0.3 Hz, control field 1 vs. shift field 1). Consistent with the observation during cue duplication, the amplitude of newly generated fields was not significantly different from the original one (averaged spiking rate: 5.0±1.0 Hz vs. 4.1±0.9 Hz, control field 1 vs. shift field 2 in Fig. 4f).

Together, our results demonstrate that CA1 place fields are strongly driven by sensory cues, independent of the cue’s absolute position within the environment, and that CA3 provides strong cue-driven input to CA1.

**Figure 4.**
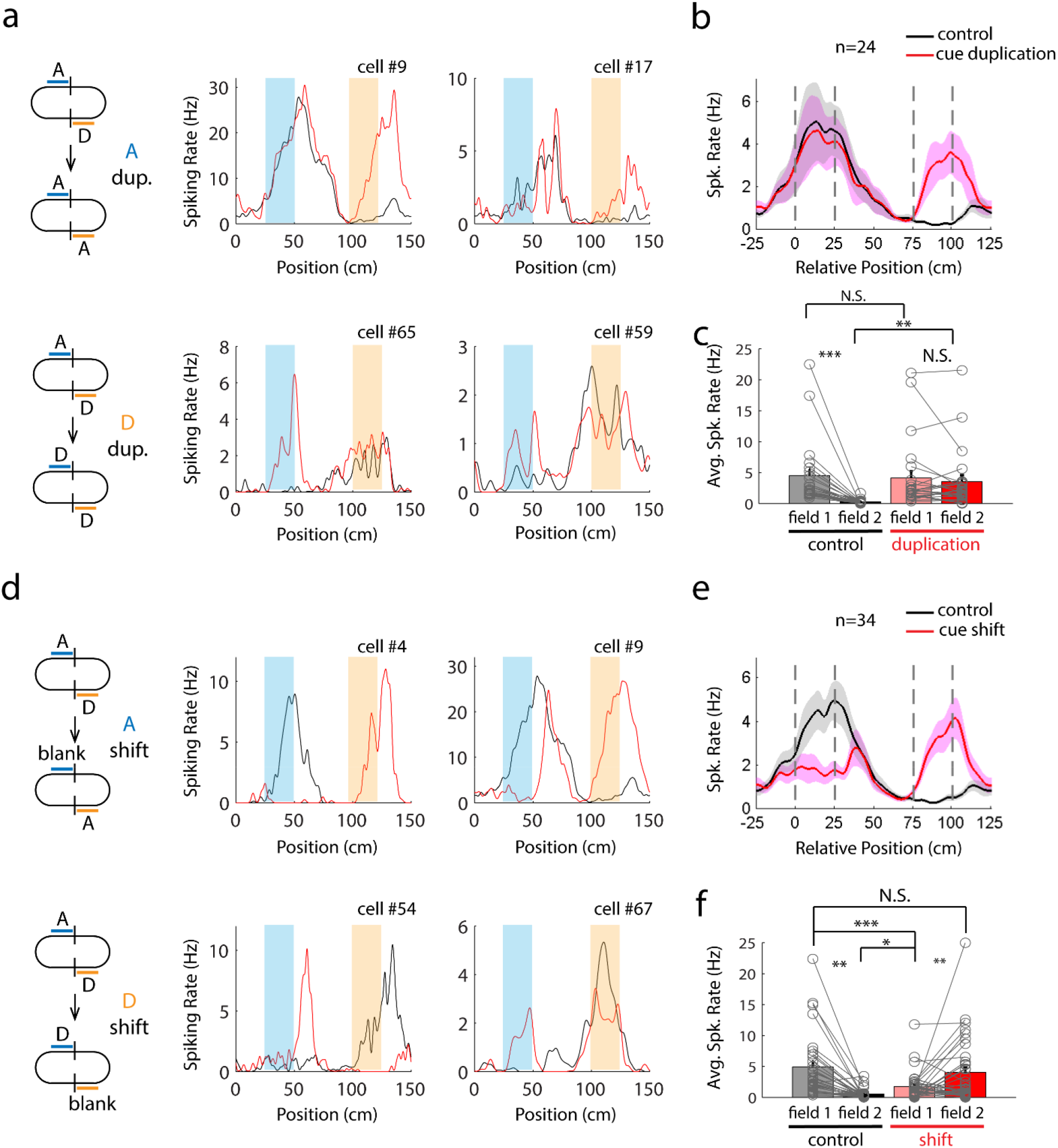
CA3 place cells followed sensory cues. **(a)** Place fields of example cells under the control condition (black) and cue duplication (red). **(b)** Spiking rates were aligned by the beginning of manipulated cues. Dashed grey lines marked the cue’s original location (0 to 25cm) and the new location where it was duplicated (75 to 100cm). Solid curves and shaded areas depict mean±s.e.m. (black for control, red for duplication, n=24 cells from 3 animals). (c) Statistics on spiking rates averaged within the original (0-25cm in b) and new (75-100cm in b) locations. Color convention is the same as in previous figures. P=3.577e-4 for control field 1 vs. control field 2, 0.395 for control field 1 vs. duplication field 1, 0.0021 for control field 2 vs. duplication field 2, and 0.341 for duplication field 1 vs. duplication field 2. **(d)-(f)** Example cells, averaged population data and statistics of CA3 place fields during cue shift. The same color conventions as in (a) to (c). P=7.504e-6 for control filed 1 vs. control filed 2, 0.0013 for control filed 1 vs. shift filed 2, 0.0166 for control filed 2 vs. shift filed 2, 0.0067 for shift filed 1 vs. shift filed 2 and 0.0767 for control filed 1 vs. shift filed 2 (n=34 cells from 3 animals, paired student t-test).

### Context-dependent sensory encoding in CA1 and CA3

After establishing that place-cell activity was tightly controlled by sensory cues, both in CA1 and CA3, we sought to test the impact of the environmental context. It has been shown that the spatial representation of hippocampal place cells completely remaps in different environments with partially overlapping sensory content; such remapping is thought to decorrelate similar inputs, thus performing “pattern separation”^11,4,25^. It is unknown, however, whether the hippocampus can also differentiate the identical sensory cue in two environments when the cue is completely separate from its environmental context (i.e. other cues).

To test for context dependency, mice were pre-trained to run in two different tracks, including the oval track described above and a triangular track (Supplementary Fig. 1c). Different sets of visual patterns were displayed along the two tracks, except for one shared cue (cue A, vertical lines). Virtual curtains were placed behind cue A on both tracks for effective isolation. Place fields in CA1 were established in the oval track through BTSP, as described above, and the virtual environment was then switched to the triangular maze. Mice were “teleported” to the triangular maze using a starting point that did not contain cue A, so that the animals could easily recognize the context change. Animals typically spun the spherical treadmill differently since the oval and triangular tracks involved turns with different angles. Animals sometimes stuck at the corner of sharp turns when they first experienced the triangular track, but learned to make turns smoothly on both tracks during the training (it should be noted though that the animal’s locomotion speed at the cue location was not different on the two tracks, as shown in Supplementary Fig. 2).

The majority of our recorded cells (n=7/8) were silent following the track change (see Fig. 5a for an example), and the only non-silent cell actually had a significant firing field that was located some distance away from cue A in the triangular maze (Fig. 5b). More strikingly, CA1 cells as a population not only lost their output (i.e., spiking fields) (Fig. 5e and 5h), they also completely lost their potentiated excitatory inputs on the triangular track. This is quantified by the massive decrease in the subthreshold ramp depolarization and theta oscillation (Fig. 5c-f; ramp: 10.7±1.4 mV vs. 2.1±0.5 mV; theta: 2.6±0.2 mV vs. 1.4±0.1 mV; rate: 16.0±3.6 Hz vs. 0.5±0.3 Hz; oval vs. triangular tracks). The differential activity cannot be explained by running speeds on the two tracks, which were similar (Supplementary Fig. 2c). These results demonstrate that the encoding of sensory cues in CA1 is strongly context-dependent, possibly due to a remapping of excitatory inputs from upstream regions.

**Figure 5.**
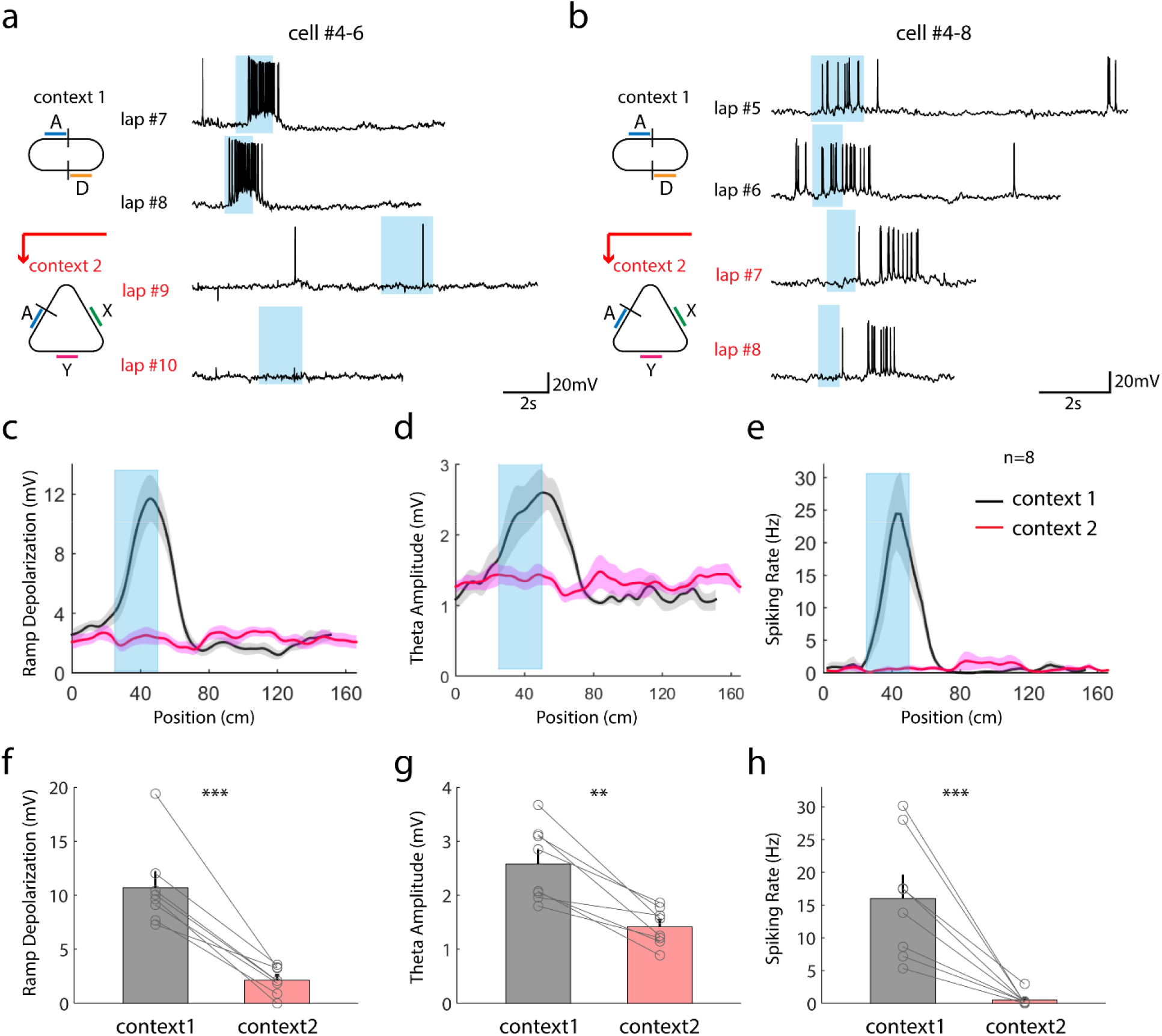
CA1 cells responded differentially to the same cue in distinct contexts. **(a)** An example cell with a robust place field on the oval track but silent on the triangular track. **(b)** An example cell with a robust place fields on both the oval and triangular tracks but at distinct locations. **(c)** Position-variant ramp depolarization on the oval (black) and triangular (red) tracks (n=8 cells from 6 animals). As in previous figures, light blue shadings mark the location of cue A shared on both tracks. Ramp depolarizations on two tracks were plotted separately because the two tracks had different lengths. **(d)** Statistical comparison between peak ramp depolarizations at cue A’s locations on the two tracks. P=6.498e-4 for oval track (context 1, grey) vs. triangular track (context 2, pink). **(e)** and **(f)** Position-variant theta amplitudes on the oval and triangular tracks and its statistical comparison. Color conventions are the same as in (c) and (d). P=0.0089 for oval vs. triangular track. **(g)** and **(h)** Position-variant spiking rates on the oval and triangular tracks and its statistical comparison. Color conventions are the same as in (c) and (d). P=1.012e-4 for oval track vs. triangular track. Paired student t-test was conducted in all analyses. Statistics labels are the same as in previous figures.

To test the hypothesis that the context dependence in CA1 is inherited from its primary driving input, we performed extracellular recordings in CA3 during the context switch. We selectively analyzed cells that had place fields overlapping the common cue shared by both environments. Among the 16 cells with place fields around the shared cue on the oval track, 10 of them became nearly silent on the triangular track (cell#2-2 and 2-9, Fig. 6a), 2 of them switched their field locations (cell #2-7) and the other 4 maintained their place fields with reduced rates (cell #2-1). As a result, the switch from the oval to the triangular track dramatically decreased CA3 place field firing at the population level (Fig. 6b, c; averaged spiking rate: 6.4±1.3 Hz vs. 1.7±0.6 Hz, oval vs. triangular tracks). Consistent with our observations of CA1 subthreshold activities, these results demonstrate that cue-driven place fields in CA3 are context-dependent.

**Figure 6.**
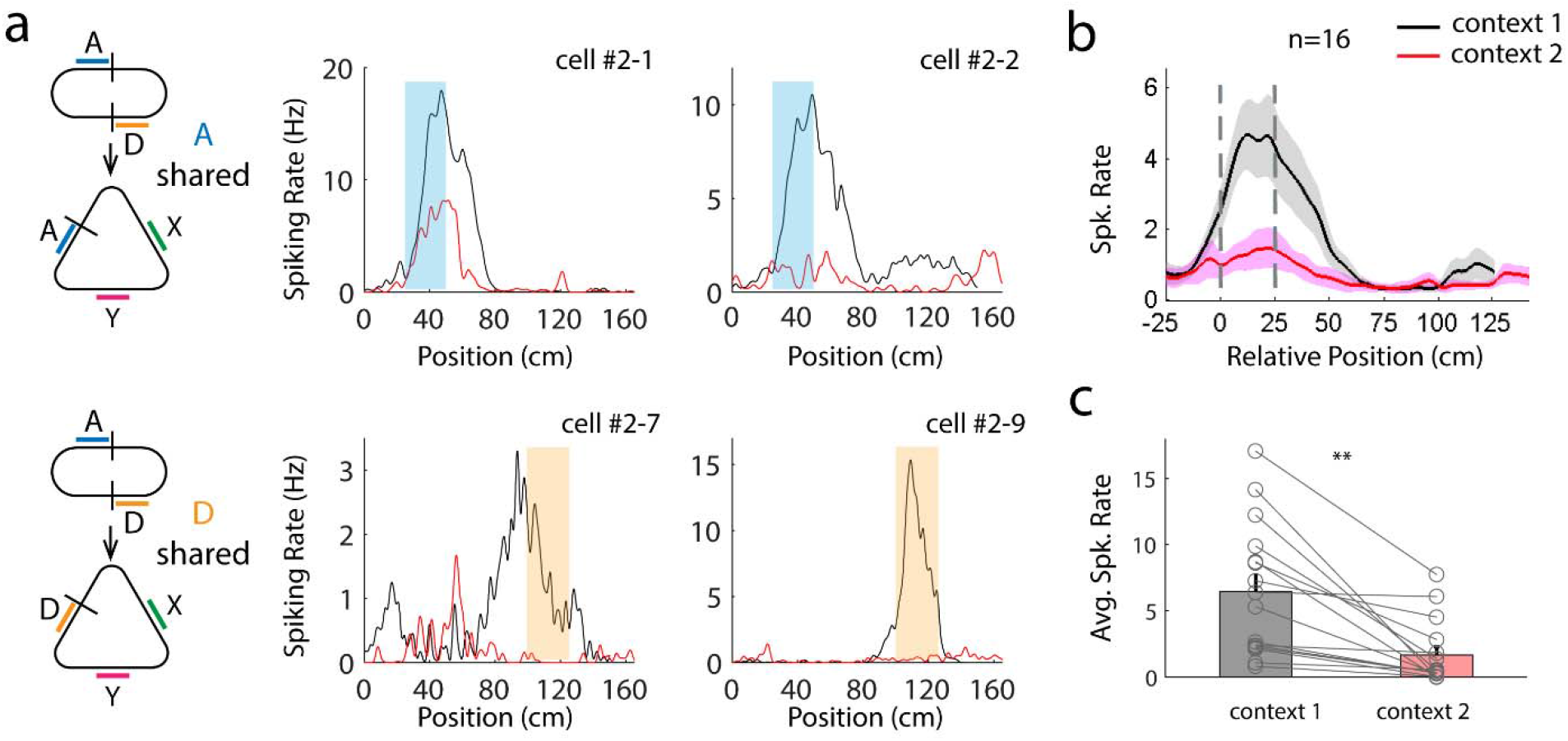
Context-dependent cue encoding in CA3 place cells. **(a)** to **(c)** Example cells, averaged population data and statistics of CA3 place fields on oval (black) and triangular (red) tracks. P=0.0013 for context 1 (oval track) vs. context 2 (triangular track), paired student t-test. Statistics labels are the same as in previous figures.

## Discussion

The results presented here provide insight into three important aspects of the encoding of sensory cues by hippocampal place cells: (1) Within a given environment, place cells can be strongly driven by visual cues inside place fields, independent of the cue’s position. (2) Visual cues evoke dramatically different responses in distinct environments. (3) The gating of visually driven place-cell firing by environmental context is observed in both CA3 and CA1, suggesting that it is generated in CA3 and/or upstream areas and inherited by CA1 pyramidal neurons via their strong input from CA3.

Mice clearly use sensory cues to navigate. For example, animals can orient themselves toward a remembered location purely relying on visual cues in the Morris water maze task^32, 33^. Here, we show that visual cues are not only necessary, but also sufficient for the formation of place-cell firing in cue-enriched virtual reality environments. Furthermore, our findings resolve some uncertainty about the relative contribution of immediate visual cues inside place fields, as opposed to cues that the animal sees elsewhere. In several previous studies, when all sensory landmarks were rotated coherently, the vast majority of place fields followed the rotation of sensory landmarks^3, 4, 34^. Although these studies provide strong evidence that sensory cues are critical in eliciting place cell firing, they do not distinguish the relative contributions of different cues. In other studies, when a subset of landmarks was moved relative to the others, a small population of place fields followed the moved cues (characterized as ‘landmark vector cells’), but this type of manipulation was only partially effective on changing place fields^9, 10, 35^. Consistent with these previous results, we found that place cells fired at lower rates when individual cues were moved but other adjacent cues were visible simultaneously. When individual cues were visually isolated, however, place-cell firing reliably followed cue movement within one environment. The virtual reality system allows us to efficiently control and isolate visual cues, through the use of virtual curtains. Combining it with the BTSP-mediated place field induction at defined locations, we establish the dominance of visual cues inside place fields in driving place-cell firing.

Place-cell responses to driver cues were modulated in an absolute manner by a change in the environmental context. Cues seen prior to the driver cue likely participate in signaling the change of environment, but we cannot rule out the possibility that mice determined the transition using other clues, such as the sharper turn angle in the triangular environment, which affected the animal’s movement during turning. Regardless, the gating of place-cell response to driver cues by environmental context was striking, as they not only stopped firing, but also showed no indication of synaptic depolarization associated with the driver cue. The strong contextual gating of place-cell firing is consistent with the notion that hippocampal place cells encode more complex cognitive variables than classically described sensory neurons.

The synaptic mechanism of context dependence in CA1 remains largely unknown. Multiple excitatory pathways, including synapses from CA3, CA2, EC, and thalamus (nucleus reuniens), converge onto CA1 pyramidal cells. One popular hypothesis is that global contextual information and local sensory cues may be channeled into CA1 through distinct pathways. Several studies have suggested that CA3 carries memory-related information that may serve to distinguish contexts, while sensory information is directly delivered by the cortical input from EC^13, 14, 36, 37^ especially the lateral entorhinal cortex (LEC)^38, 39^. In addition, a recent study reported that most CA3 cells did not encode sensory landmarks, and thus argued that the sensory information in CA1 might originate from other sources, such as CA2^16^.

Our intracellular recordings shed light onto whether signals from one sensory cue and its context are first combined in the CA1 subregion, or more upstream areas. If CA1 were indeed the first stage to integrate signals from local cues and context, the subthreshold V_m_ of CA1 cells would be expected to show a smaller ramp or a more hyperpolarized baseline after the context change. However, our results revealed that the primary cue did not evoke a measurable depolarization of V_m_ in a new context. These data suggest that after a context switch, the vast majority of driver cue-associated excitatory input to CA1, which was potentiated by BTSP during induction of the place field in the original context, is no longer active in the new context.

Our data also offer some insight into the mechanisms by which synaptic depolarization in CA1 is gated by context. Specifically, our CA3 recordings showed that most CA3 place cells followed sensory cues and remapped in different environments, with very little spiking observed in response to the driver cue in the new environment. Thus, in the new context, the driver cue provides very little synaptic drive from CA3 to CA1. However, postsynaptic CA1 cells may integrate CA3 inputs in a non-linear manner^40–42^, thus enhancing changes in input activity. As a result, small amounts of input from CA3 may not be detected as significant synaptic depolarization in CA1. Furthermore, recruitment of synaptic pathways other than CA3 by BTSP, which may be differently affected by cues/contexts complicates this issue. In fact, plateau potentials and dendritic spikes in CA1 cells can potentiate coactive EC inputs^43–45^, although the time window of pre-/post-synaptic activation required for such plasticity has not yet been well characterized. How these secondary excitatory inputs to CA1 are integrated with CA3 inputs to drive and modulate place-cell firing should continue to receive attention, not only for their role in synaptic integration, but also for the potential implications for synaptic and circuit plasticity^45–47^. Nevertheless, our data reinforce the idea that CA3 provides the main spatially tuned excitatory input to CA1 and that potentiation of these synaptic weights through BTSP provides the V_m_ depolarization that drives place field firing in CA1, and that CA1 place cells undergo context-dependent remapping because they inherit a completely different input pattern from CA3.

Anatomical and in-vitro physiological studies have identified that CA3 provides numerically the largest^27, 48^ and functionally the strongest ^28–30^ excitatory synapses to CA1. Paradoxically, in-vivo recordings have indicated that basic CA1 place field properties were nearly unaffected following chronic CA3 lesion/silencing^17, 31, 37^, challenging the idea that CA3 is the major excitatory driver of CA1. Recently, however, a study using acute silencing of CA3 revealed a dominant CA3 input to CA1^49^. Our results support the primary role of CA3 in driving CA1 place cells.

The notion that the hippocampal representation of space is context dependent is central to the concepts of the cognitive map, as well as its putative roles in episodic memory and even human thought^1, 2, 50–52^. Beside environmental contexts, other factors are known to influence place-cell firing. For instance, animals may rely on non-sensory landmark dependent strategies to navigate in cue-impoverished zones (e.g. path integration^34, 53^). In addition, behavioral trajectory in a task involving alternating turns can modulate place-cell firing^15, 54, 55^, as can the introduction of rewards at consistent locations^20^. In future studies, it will be interesting to investigate whether similar synaptic mechanisms mediate contextual gating under these scenarios. Understanding the circuit, synaptic, and biophysical mechanisms by which these and other factors determine the hippocampal code for space is a central challenge for mechanistic cognitive neuroscience. Whole-cell patch-clamp recording in virtual reality is a powerful approach for making progress toward this goal.

## Methods

### Animal surgeries

Male C57Bl/6 mice (post-natal 60-120 days, Charles River) were used in all experiments. Animals were anesthetized by isoflurane (3-4% for induction and 1.5-2% for maintenance) and mounted on a stereotaxic (Kopf) with ear bars. For CA1 intracellular recordings, the animal’s skull was first exposed, cleaned and covered with an instant adhesive. Customized titanium headbars were implanted on the animal’s skull using dental acrylic (Ortho-Jet, Lang Dental Manufacturing Co.), as described before^25^. After recovering from surgeries for about 1 week, mice were transferred to the water restriction schedule, receiving 1.5ml water each day.

For intracellular CA1 recordings, a small craniotomy (~200um in diameter) was made over the right hippocampus (2.0mm posterior and 1.7mm lateral from Bregma) after training. A second craniotomy was made 1.4mm behind the first one, through which local field potential (LFP) was monitored in the hippocampus through a second pipette. Recordings started the next day after craniotomy surgeries, and continued for 2-4 days. Craniotomy windows were sealed by silicone elastomer (Kwik Cast, World Precision Instrument) between each recording day.

For extracellular recordings in CA3, six-shank H64LP silicon probes (NeuroNexus) were mounted on the customized titanium micro-drives using super glue and Ortho-Jet cement.

Male C57Bl/6 mice were anesthetized by isoflurane. A titanium head plate was first implanted on the animal’s skull using the same approach as described above. A small hole was drilled on top of the cerebellum (6.0mm posterior and 2.8mm lateral from Bregma). A ground screw (~0.7mm diameter, with a thin copper wire pre-soldered on its cap) was inserted into cerebellar cortex through this hole, and then mounted on the skull with Ortho-Jet cement. A 1.5mm diameter craniotomy was open, centered at 2.35mm posterior and 2.55mm lateral from Bregma. The micro-drive, with the probe on it, was mounted on a manual micromanipulator (Narishige) through a customized adapter. The plane of probe shanks was aligned 45° relative to the midline to match the longitudinal axis of hippocampus, and carefully centered in the craniotomy window. The probe was then tilted by 10-12° vertically to the anterior direction. This configuration was to increase the chance that multiple shanks could get into the CA3 pyramidal layer together, thus maximize the number of simultaneously recorded cells. The probe was slowly inserted into the brain with the micromanipulator, 1.1mm below the pia. Ortho-Jet cement was used to mount the micro-drive onto the skull. In addition, a circular well was made around the micro-drive and probe using Ortho-Jet cement. After complete drying of the cement, the micro-drive was released from its holder. The attached electronic circuit was glued and cemented on the outside surface of the cement well. Silicone gel (4680 silicone gel kit, Dow Chemical) was poured into the well to cover the craniotomy and probe shanks. Finally, a copper mash cone was made around the whole implant to protect the recording apparatus and shield it from noise. Both the ground wire from the probe and the one attached to the ground screw were soldered on the copper mash.

All experiments were performed according to protocols approved by Janelia Research Campus Institutional Animal Care and Use Committee.

### Virtual reality system

Behavioral training started 1 week after the water restriction schedule began. Animals were head-fixed on a stainless steel headbar holder, and trained to run on top of a 40cm diameter hollow foam ball. There thin-bezel monitors (463UN, NEC) were placed in front of the spherical treadmill, which covered 216° horizontally and 91° vertically (27° below its eyes and 64° above) of the animal’s visual field. The ball’s movement was tracked by two customized high-speed infrared cameras^19^. Optical flow at all three rotation axes (pitch, roll and yaw) were calculated by an optical mouse sensor chip (ADNS-6090, Avago).

3-D virtual environments were first created in Blender, an open-source animation software. Sizes and contrasts of these visual cues were designed to optimize potential visual responses based on our knowledge of the mouse visual system. Calculated from the center of the track, the spatial frequency of gratings ranged from 0.02 to 0.04. The size of each circular dot was 10° in view angle. In addition, stripes were painted on the floor at turning locations to facilitate the animal’s perception of rotation. The virtual scene was rendered on monitors at 60Hz by Janelia’s open-source virtual reality platform Jovian (https://github.com/JaneliaSciComp/-CohenBolstadLee_eLife2017). All displays were black and white. The ‘sky’, i.e. empty space in the scene, was rendered as white. The roll/pitch ratio of the spherical treadmill’s movement, determined by the animal’s body angle during running, was used to calculate its head direction in virtual environments. When the animal hit a wall, Jovian simulated the animal sliding along it, by projecting the animal’s locomotion velocity onto the wall’s direction. To improve the smoothness of the animal’s turning, an automatic heading correction mechanism was implemented as described before^19^.

Animals were rewarded with 10% sucrose water controlled by a solenoid valve (EV-2-12, Clippard). The animal’s licking was monitored by an optical beam breaker sensor (FX-301H, with FT-V22 optic fibers, Panasonic) placed on both sides of the lick port. The first 2-3 days of training included two short sessions (20-30min each, with 3-5h in between) each day. Animals were then trained one session per day for 1-2 weeks, with the session duration gradually increased from 30min to 60-90min. At the end of training, animals typically ran 100-200 laps within one session.

Virtual cue manipulations were done by switching virtual tracks in Jovian. All monitors were blacked out for 4-5s between switches, during which animals did not get any reward regardless of their behaviors. In experiments where cues were manipulated within the same track, the last 3-4 days of training contained 5-6 epochs each day, separated by these short black-out gaps. At the end of the training, animals adapted well to the black-out without any struggling. The same track (no cue manipulation) was loaded after each gap. As a result, in all experiments with cue manipulations within the same environment, animals were not exposed to manipulated cues during training. However, our dataset includes some sessions in which the animal already experienced cue manipulations in earlier recordings. These cells were pooled together with others, in which the animals saw the cue manipulation in the first time, since there is no significant difference between the two groups. In experiments where both oval and triangular tracks were tested, animals were initially trained on the oval track. In the last 3-4 days, animals experienced both oval and triangular tracks in different epochs in a random order.

Reward delivery and recording of the animal’s behavior were implemented by a micro-controller (chipKit Max32, chipKit) with customized firmware developed in MPIDE (chipKit).

### Intracellular recordings in CA1

Whole-cell recordings were made with glass pipettes (9-12MOhm), using a standard blind patching technique^25^. Pipettes were filled with internal solution containing (in mM) 134 potassium gluconate, 6 KCl, 10 HEPES, 4 NaCl, 0.3 Mg-GTP, 4 Mg-ATP, and 14 Tris-phosphocreatine. In some recordings in which recorded cells were reconstructed, 1% biocytin (Sigma) was added. Pipettes were inserted into the brain with a micromanipulator (Junior 4 axen InVivo Unit, Luigs and Neumann). High positive pressure (~10psi) was used during the penetration of dura to avoid potential clogging. The pressure was reduced to 5-6psi when the pipette was in cortex (500-700μm under pia). The pipette was advanced into the hippocampus, and the pressure was further reduced to around 0.3 psi before we started to search for cells. Standard seal and break-in procedures followed after the detection of reproducible impedance increase^25^. A second pipette, filled with 0.9% saline, was inserted at 45° into the hippocampus using a hydrolic micro-manipulator (Narishige) through the posterior craniotomy to monitor LFP and help stabilize the recording mechanically. During recordings, the craniotomy was covered with 0.9% saline. Membrane potential was recorded in current-clamp mode using a Dagan BVC-700A amplifier, and digitized with a National Instrument board (USB 6343). Data acquisition was done with Wavesurfer, a MATLAB-based electrophysiology platform developed at the Janelia Research Campus (http://wavesurfer.janelia.org/). Bridge balance was carefully adjusted to fully compensate the series resistance (Rs). Recordings with Rs>100MΩ were excluded from analyses. During the pipette placement and insertion, the VR display was off and the animal’s eyes were protected from the surgical lamp by a black visor. Junction potential was not corrected in all reported numbers in this study.

To synchronize behavioral and electrophysiology recordings, 1-Hz 10ms TTL pulses were generated by a pulse generator (Master-9, AMPI), and recorded in both behavior and electrophysiology data acquisition systems.

### Extracellular recordings in CA3

Mice recovered for 1 week from surgeries before being transferred to the water restriction schedule (1.5ml water per day). Training in the virtual reality system started 4-7 days after the beginning of water restriction as described above for intracellular recordings. During the animal’s training period, the probe was slowly advanced toward the CA3 region with the micro-drive (12.5μm at a time; at least 1h interval between consecutive movements). Signals were filtered, amplified and digitized by Amplex (KJE-1001) to check for characteristic hippocampal sharp-wave and theta activities. Signals were acquired and visualized with NeuroScope. CA3a/b zones were targeted to avoid potential confound from dentate granule and mossy cells. Most probes were located within or close to the pyramidal layer of CA3a, which was characterized by strong LFP theta oscillation and large amplitude spikes across a long range of recording sites (>200μm), due to the thick and curved CA3a pyramidal layer. Consistently, sharp wave signs did not reverse across this range. For a few shanks, spikes were detected at shallow depth (<1300μm) and sharp waves/ripples quickly reversed within ~50μm, characteristic for CA1/CA2 localization. In this cases, probes were further advanced until large spikes were detected again in CA3b (typically after 500-600μm). Probe locations in all animals were verified by histology after all recording sessions were done.

Recordings began once high-quality spikes were detected on some probe shanks. Two cues (A and D, see Fig. 1b) have been manipulated throughout the current study. One recording session, with the manipulation of one cue, was performed each day (see Supplementary Fig. 5 for details). A recording session with the other cue’s manipulation was performed on the second day without any movement of the probe. Although we optimized our probe mounting angles, shanks of one probe could still reach the CA3 pyramidal layer at different depths, due to its curved geometry. The probe was thus further advanced by 100μm within 2-3 days, and recordings were conducted again. This process was repeated for multiple rounds until no spikes appeared on new recording sites. The current study included 2 rounds of recordings in animal #1, 3 rounds in animal #2 and 1 round for animal #3.

Averaged voltage trace from reference channels was first subtracted from signal recorded on each site to remove movement artifacts. Spikes were filtered and detected with NDManager as described before^56^. Single units were then isolated in KlustaKwik by clustering spikes based on principle components of their waveforms (Supplementary Fig. 5). The clustering was first done automatically and then manually curated. Autocorrelograms of spike timing were carefully validated for each unit to make sure no spikes fell into the window of typical refractory period (2ms).

### Histology

To reconstruct intracellularly recorded cells, mice were euthanized and perfused with first PBS and then 4% paraformaldehyde (PFA) 2-3 hours after the recording. The animal’s brain was dissected out and further fixed in 4% PFA for overnight. 150μm thick coronal sections were cut using a vibratome (Leica). Brain slices were first penetrated in 0.1% Triton-X (Sigma) for 1h, and then transferred into 1:800 diluted Alexa-488 conjugated streptavidin (Molecular Probes) for 2-4h. Slices were next rinsed in PBS for 3 times (5min each), and mounted on coated glass slides with mounting medium containing DAPI (Vectorshield, Vector Laboratories).

To localize silicon probe tracks in CA3 recordings, mice were euthanized and perfused with 4% PFA after the all recording sessions were done. 50μm thick coronal sections were cut using the Leica vibratome. Brain slices were mounted with mounting medium containing DAPI as described before.

### Data Analysis

Animals ran on narrow virtual tracks in this study. Raw running speed of the animal was calculated at each sampling point as the distance it moved from the last point in the virtual environment divided by the sample time interval. The speed was smoothened by a Gaussian kernel with a standard deviation of 50ms before further analysis. The animal’s position is linearized to one dimension by first projecting the animal’s 2-dimensional coordinates onto the nearest point along the track’s midline; and then the path distance was calculated between this point and an arbitrarily chosen ‘lap start point’. It should be noted that the ‘lap start point’ is purely for data presentation purposes. The animal did not experience any discontinuity at this position.

Hippocampus operates in distinct modes depending on the animal’s behavioral states. Throughout this study, we only included data during the running period (speed>2cm/s) in our analyses.

To analyze membrane potential (Vm) from intracellular recordings, electrophysiology and behavioral recordings were aligned through 1Hz synchronization pulses. Behavioral data was interpolated to match the sampling rate of electrophysiology recording.

We first corrected a slow artificial drift of Vm readings during each recording session, primarily due to the change of solution level covering the animal’s craniotomy. Evaporation of the solution caused parallel drifts for both the patch and LFP pipettes. Since the physiological fluctuation of LFP is much faster than such drift, the mean LFP was calculated for each lap, and used to offset the Vm recording.

Action potentials were detected by dVm/dt>40mV/ms. In a small set of recordings that contained some spikelets, the threshold was increased to 60-70mV/ms, and recording traces were visually inspected to avoid inclusion of any spikelets. To analyze subthreshold activity, action potentials were first removed by a median filter with 10ms window. Ramp-Vm and theta-Vm were then extracted by bandpass filters (<2Hz for ramp and 4-10Hz for theta). Ramp depolarization was calculated by subtracting the minimal value of Vm, averaged across all control laps, from the Ramp-Vm (Fig. 2, 3, and 5, S3 and S4). Theta oscillation amplitude was calculated as the Hilbert transform of theta-Vm.

To calculate the position-variant ramp depolarization and theta amplitude, the track was divided into 0.1cm bins. Measured values were averaged among all sample points at which the animal’s positions fell into each spatial bin. The resulted spatial tuning curves were then smoothened by circular convolution with a Gaussian kernel (standard deviation: 2.5cm). To calculate the position-variant spiking rate, the track was divided into 2.5cm bins. Spiking rate in each bin was calculated as the number of action potentials divided by the total occupancy time within that spatial bin. Spiking rate curves were further smoothened by a Gaussian kernel with standard deviation of 5cm.

Statistical comparisons of ramp depolarization, theta oscillation and spiking rate (bar plots in Fig 2,3 and 5, S3 and S4) were all based on peak amplitudes. For each measurement, the position where the maximal value was located within each cue’s range (between the two dashed light blue or yellow lines) was first determined. The peak amplitude was calculated as the mean within a 10cm window center at the peak position.

To analyze extracellular recordings, electrophysiology and behavioral recordings were first aligned and interpolated in the same way as described above. Spiking rates were calculated within 0.1cm spatial bins, and then smoothened by a Gaussian kernel with 5cm standard deviation. Each recording session was comprised of one or two cue manipulation epochs interleaved with control epochs. Rate maps, under control and cue manipulation conditions, were calculated by averaging position-rate curves across all laps in different epochs, respectively.

Only cells that satisfied the following three criteria under the control condition were included in our place field analysis: 1) peak spiking rate>1Hz; 2) spatial selectivity (SS) > 0.3 and 3) Pearson’s correlation coefficient between rate maps under control epochs > 0.3. Spatial selectivity was quantified as 1-circular variance:

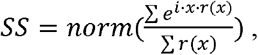

where x is the position vector converted to 0-2π scale and r(x) the spiking rate map. The selection of place cells was only based on their activities under control conditions, regardless of their responses to cue manipulations.

For further place field analyses, rate maps from multiple control epochs were averaged as the rate map for the control condition.

Place cells were sorted by the center of mass (COM) of their place fields, calculated as the spiking rate-weighted circular mean:

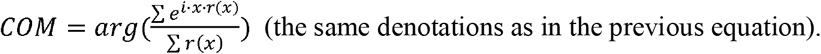

To identify place field locations in extracellular recordings (Fig. S6), spiking rate maps were fitted using the skewed von Mises equation, an equivalent of skewed Gaussian in the circular dimension:

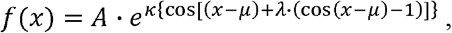

where x is the position, A the scaling factor, 1/κ the variance, μ the shift and λ the skewness factor. Circular distribution was used because of our track’s periodic nature. The peak location was determined by the position of the maximal value in the fitted curve. Positions on the fitted curve that crossed the 20% peak amplitude threshold were defined as the starts and ends of place fields.

Two-tailed t-test was conducted in all significance tests between paired samples. Bar graphs and error bars represent mean and s.e.m., respectively.

## Acknowledgement

We thank Mark Bolstad, Steven Sawtelle, Dr. Albert Lee and Dr. Jeremy Cohen for their help on the design and manufacture of virtual reality system. We thank Dr. Katie Bittner for her development of intracellular recording techniques. We thank Dr. Brett Mensh and all Magee and Spruston lab members for insightful discussions. This study is supported by the Howard Hughes Medical Institute.

## Author Contribution

XZ, NS, and JCM designed experiments; XZ performed in vivo whole-cell recordings; XZ and YW performed extracellular recordings; XZ and YW analyzed the data; XZ, NS and JCM wrote the manuscript with input from YW.

## Supplementary Figures

**Figure S1.**
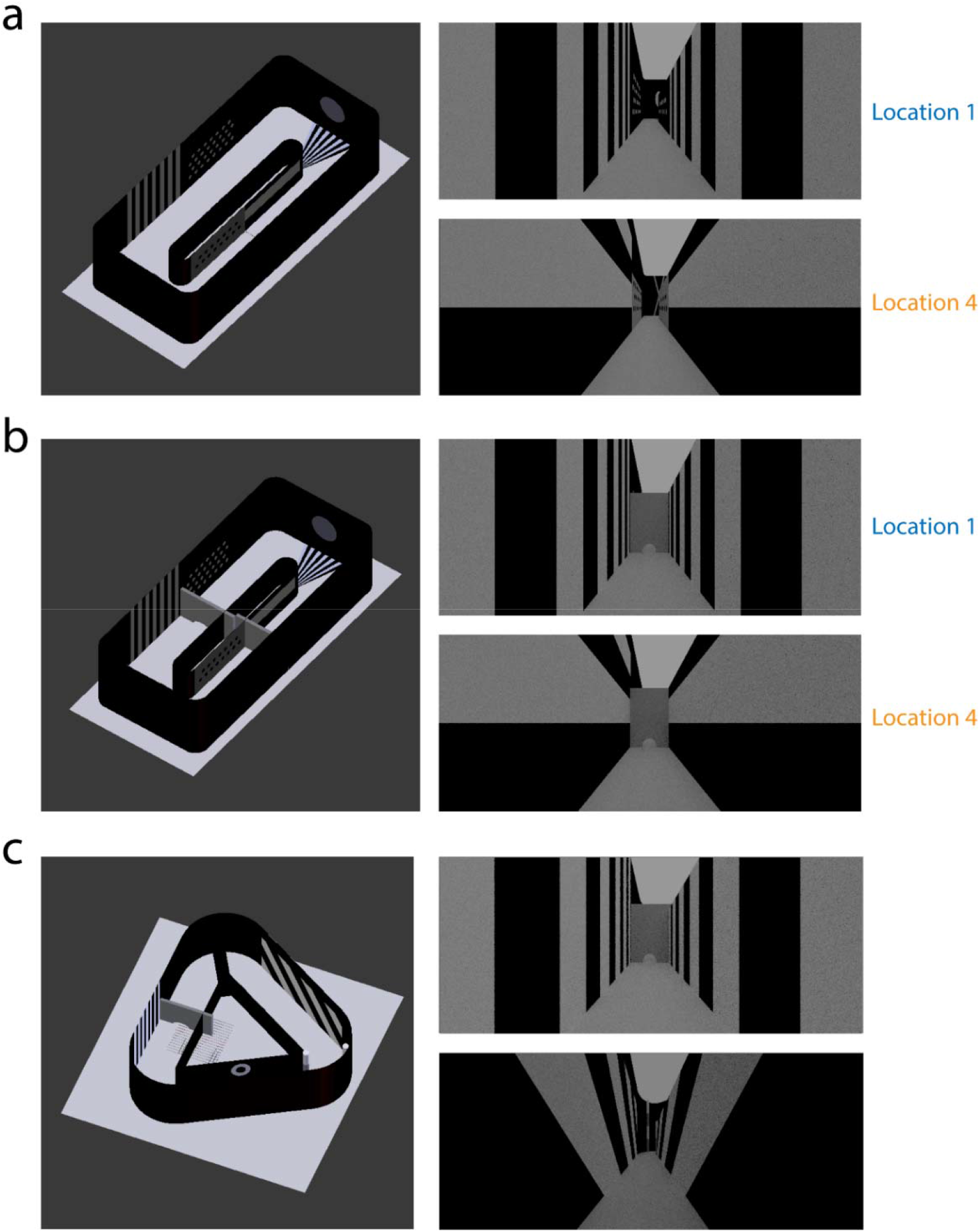
Virtual tracks with distinguishable visual cues. Oval track (a), oval track with barriers (b) and triangular track (c) used throughout this study. Besides the common cue (vertical gratings) shared on both tracks, the triangular track contained another 2 cues (45° gratings and rings) that were not present on the oval track.

**Figure S2.**
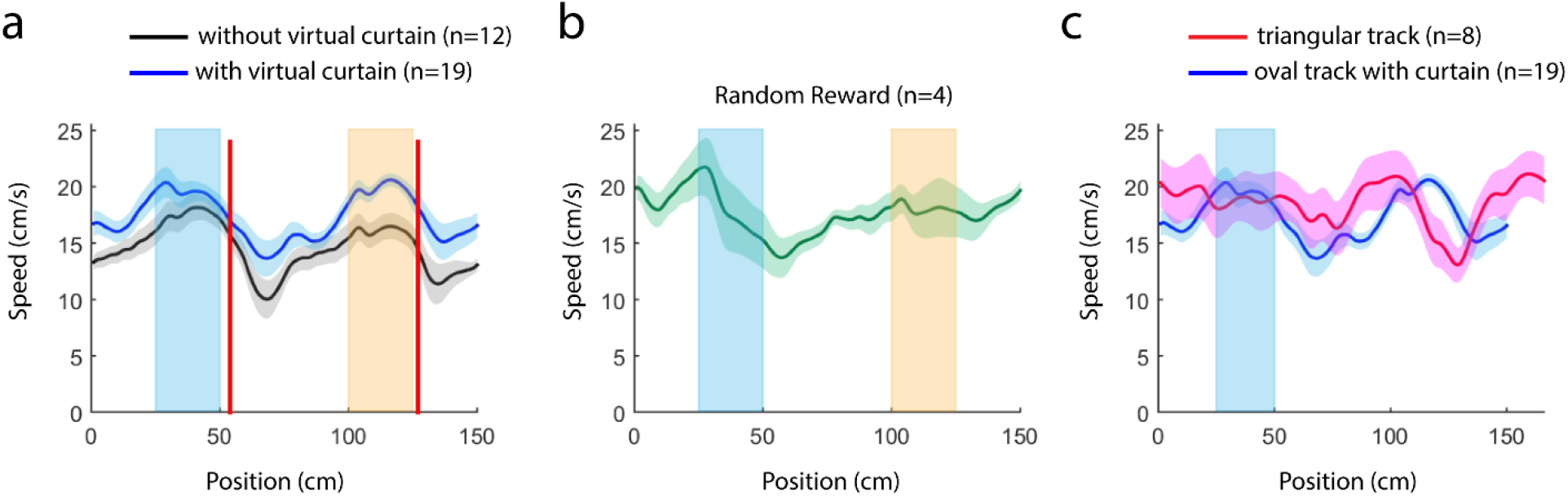
Running speeds during different behavioral paradigms. (a) Position-variant running speeds on oval tracks with (blue) or without (black) virtual curtains in the fixed reward paradigm (reward locations marked as red lines). Although the overall averaged speeds slightly differ under the two conditions due to variations among individual animals, their position dependencies are very similar, likely determined by the turning and reward locations. (b) Position-variant running speed on the oval track with random reward delivery. Dashed lines depict location 1 and 4, as in Fig. 1. (c) Position-variant running speed on the triangular track (red). Speed on the oval track with virtual curtains was plotted as comparison (blue). Shadings depict the location where the same cue was available on both tracks. In all panels, population data are presented as mean±s.e.m.

**Figure S3.**
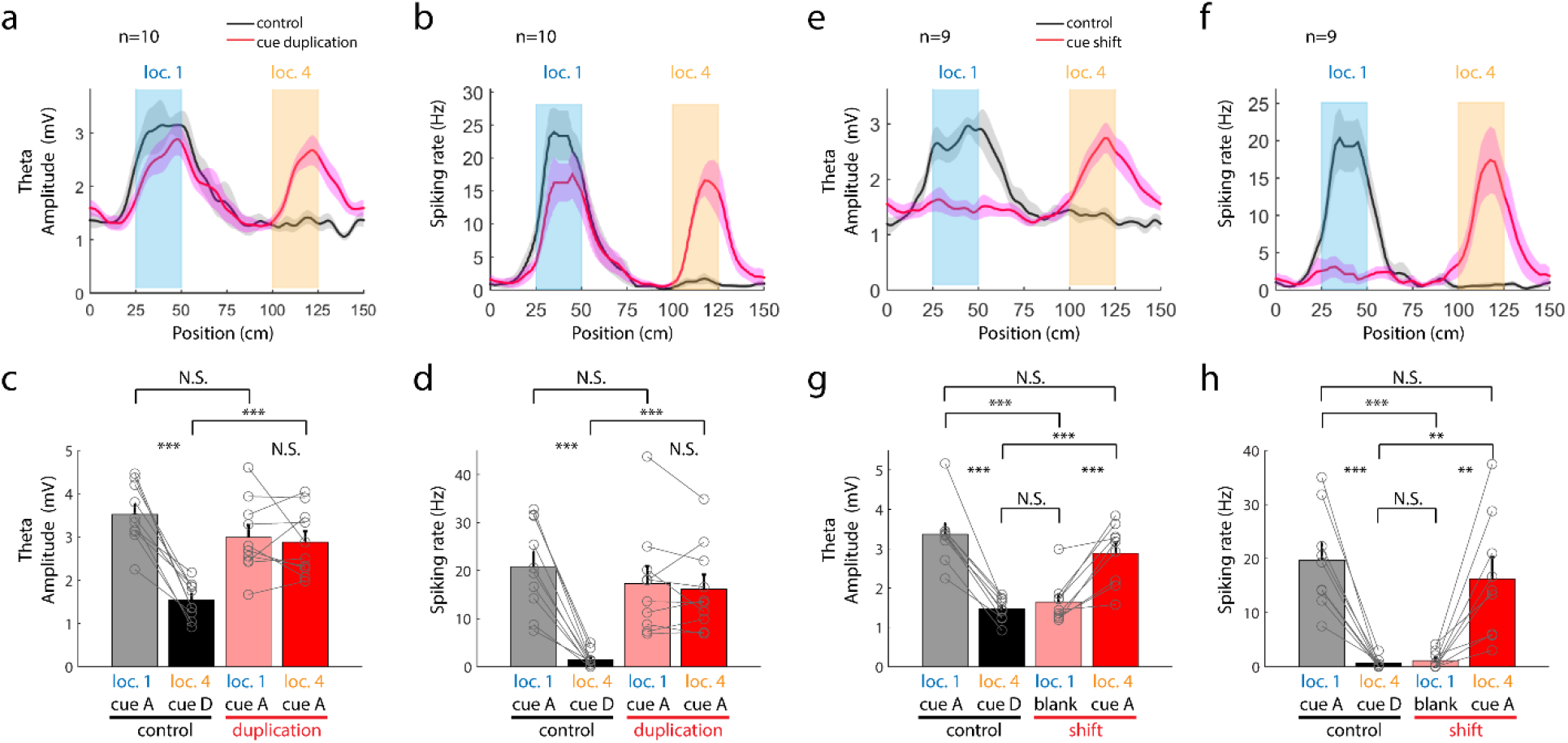
Detailed characterization of place fields during manipulations of isolated cues. **(a)-(d)** Population averages and statistics of theta amplitudes and spiking rates following the cue duplication. All data are presented as mean±s.e.m. The same color conventions as in Fig. 3. In (c): P=1.936e-5 for control loc. 1 vs. control loc. 4, 0.641 for duplication loc. 1 vs. duplication loc. 4, 0.0564 for control loc. 1 vs. duplication loc. 1, and 3.451e-4 for control loc. 4 vs. duplication loc. 4. In (d): P=3.97e-5 for control loc. 1 vs. control loc. 4, 0.5135 for duplication loc. 1 vs. duplication loc. 4, 0.1553 for control loc. 1 vs. duplication loc. 1, and 2.04e-4 for control loc. 4 vs. duplication loc. 4. **(e)-(h)** Population averages and statistics of theta amplitudes and spiking rates following the cue shift. Paired student t-test was conducted in all analyses. In (g): P=2.744e-5 for control loc. 1 vs. control loc. 4, 1.85e-4 for shift loc. 1 vs. shift loc. 4, 2.454e-4 for control loc. 1 vs. shift loc. 1, 0.0632 for control loc. 1 vs. shift loc. 4, 9.723e-4 for control loc. 4 vs. shift loc. 4, and 0.2953 for control loc. 4 vs. shift loc. 1. In (h): P=1.959e-4 for control loc. 1 vs. control loc. 4, 0.0035 for shift loc. 1 vs. shift loc. 4, 2.667e-4 for control loc. 1 vs. shift loc. 1, 0.2406 for control loc. 1 vs. shift loc. 4, 0.0034 for control loc. 4 vs. shift loc. 4, and 0.4954 for control loc. 4 vs. shift loc. 1.

**Figure S4.**
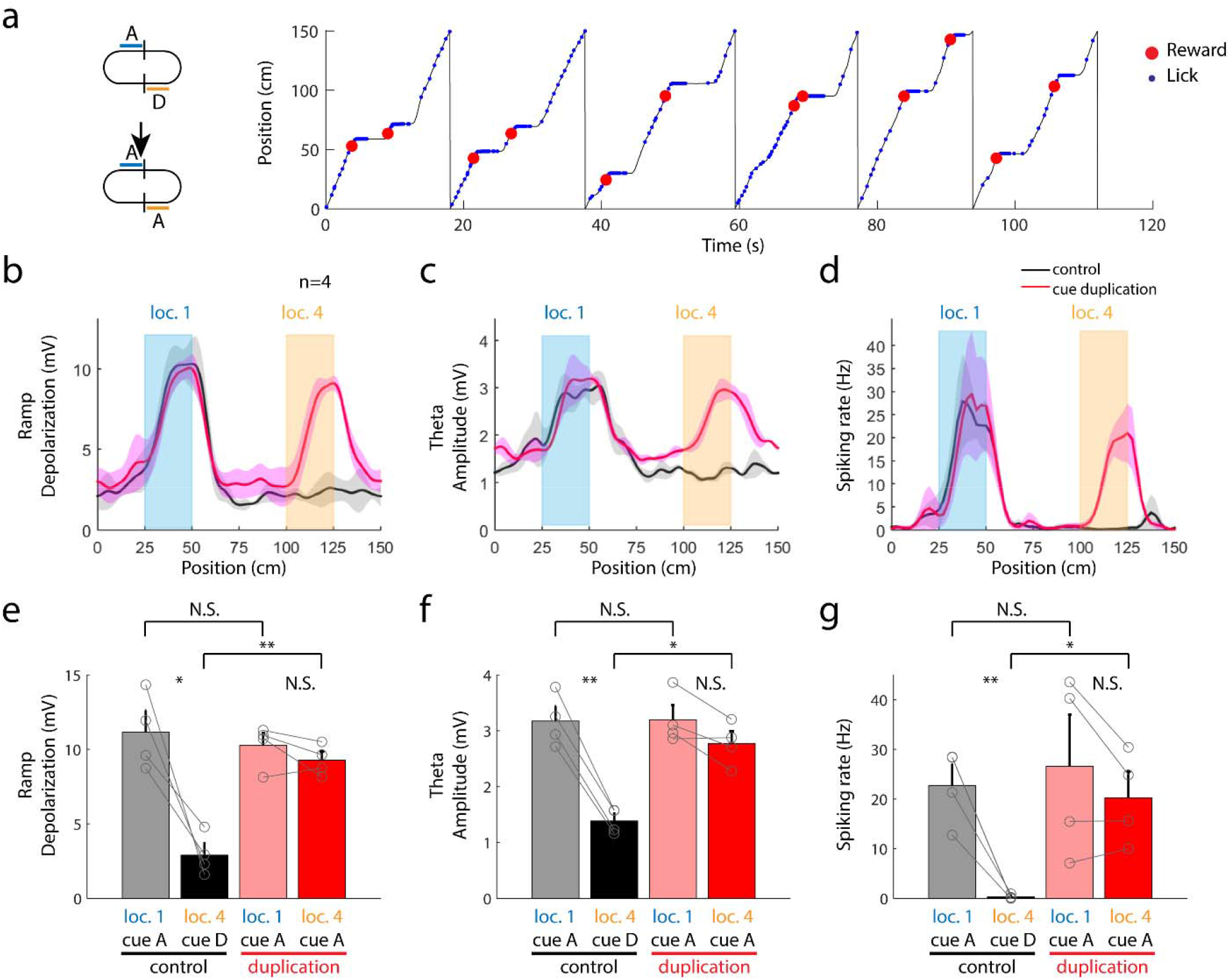
Effects of cue duplication on CA1 place fields with random reward delivery. **(a)** Behavior of an example animal. Two rewards (red) were delivered in each lap at random locations. Note that the animal licked (blue) along the entire track. (b) Population averages and statistics of ramp depolarizations, theta amplitudes and spiking rates following the cue duplication. All data are presented as mean±s.e.m. The same color conventions as in Fig. 3. Paired student t-test was conducted in all analyses. In (e): P=0.0204 for control loc. 1 vs. control loc. 4, 0.2255 for duplication loc. 1 vs. duplication loc. 4, 0.6152 for control loc. 1 vs. duplication loc. 1, 0.2318 for control loc. 1 vs. duplication loc. 4, and 0.0019 for control loc. 4 vs. duplication loc. 4. In (f): P=0.0011 for control loc. 1 vs. control loc. 4, 0.0794 for duplication loc. 1 vs. duplication loc. 4, 0.9538 for control loc. 1 vs. duplication loc. 1, 0.3426 for control loc. 1 vs. duplication loc. 4, and 0.0169 for control loc. 4 vs. duplication loc. 4. In (g): P=0.0091 for control loc. 1 vs. control loc. 4, 0.2591 for duplication loc. 1 vs. duplication loc. 4, 0.6522 for control loc. 1 vs. duplication loc. 1, 0.5469 for control loc. 1 vs. duplication loc. 4, and 0.0216 for control loc. 4 vs. duplication loc. 4.

**Figure S5.**
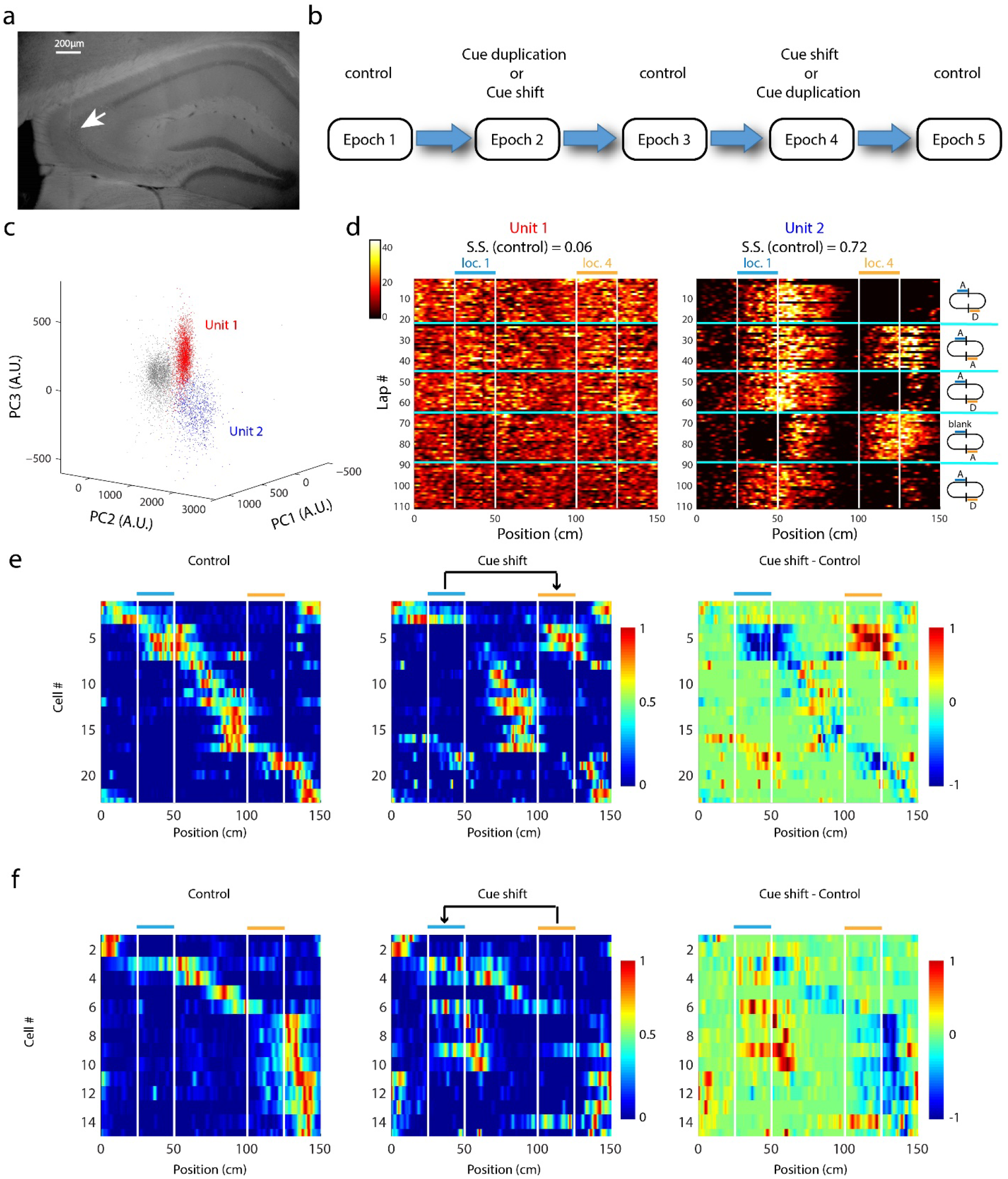
Isolation of single units from extracellular recordings in CA3. **(a)** Representative histology confirmation of extracellular recordings in CA3 using silicon probes. **(b)** Recording paradigm. Most of recording sessions (n=7/9) in this report included 5 epochs. Two cue manipulation epochs (duplication and shift) were sandwiched between control epochs. The rest 2 sessions only included the cue shift epoch and two control epochs. **(c)** Spike clustering based on principle component analysis (PCA) of spike waveforms. Red and blue dots are two isolated spike clusters, while grey dots represent all other spikes. **(d)** Color coded rate maps of the two sorted units shown in c (color scale: 0-45Hz). Unit 1 is a putative interneuron (spatial selectivity: 0.06). Unit 2 is a place cell (spatial selectivity: 0.72). Boundaries between epochs are marked by cyan lines. Vertical white lines mark the locations of cue A and B, as in Fig. 5. In this recording session, the second and fourth sessions were cue A’s duplication and shift, respectively, as labeled on the right. **(e)** and **(f)** Example sessions of simultaneously recorded place cells in CA3 during shift of cue A (e) or B (f). Spiking rate of each cell was normalized to its peak rate under the control condition. Cells were sorted based on center of mass of their firing field under the control condition.

**Figure S6.**
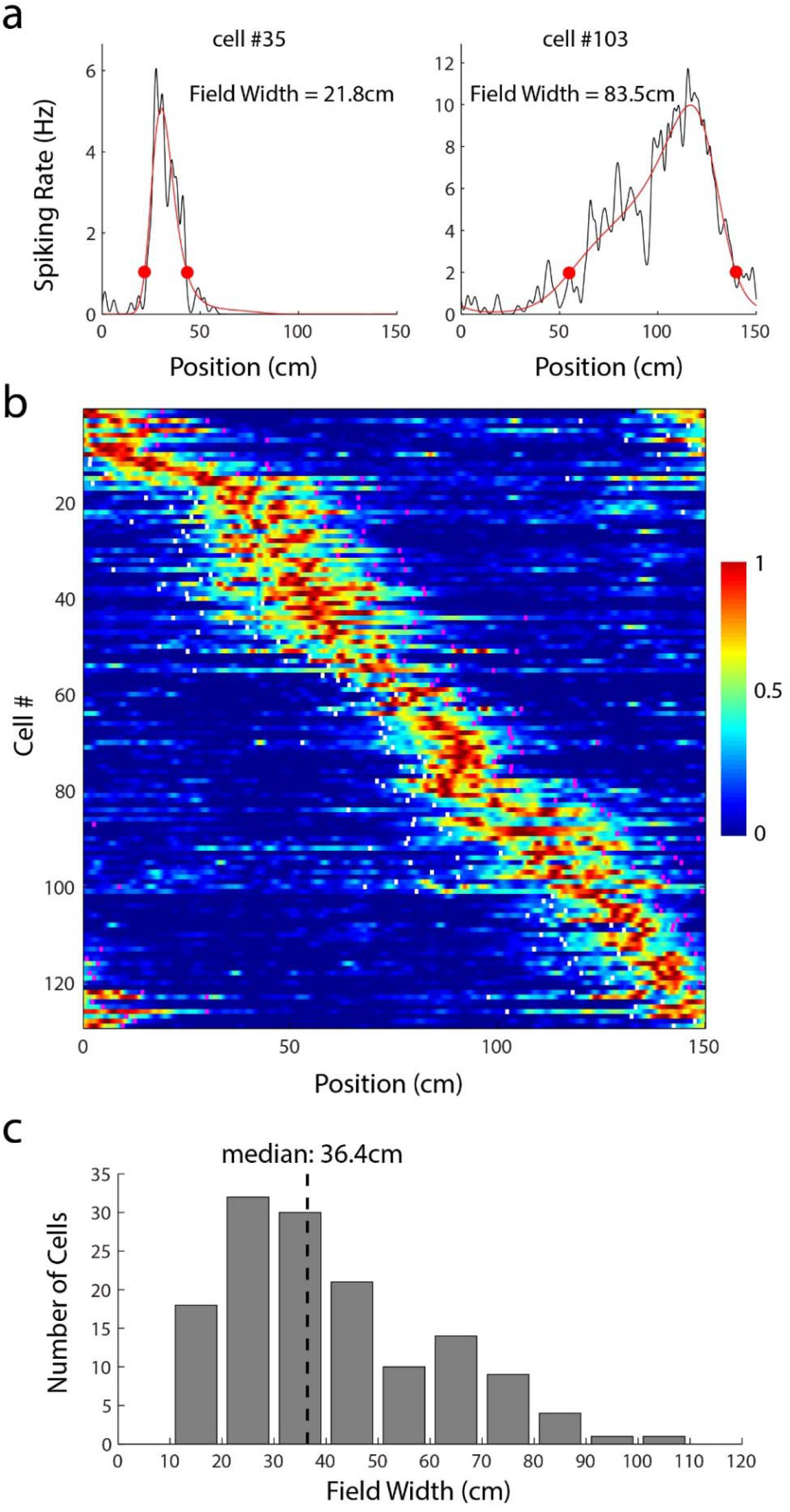
Identification of place fields in CA3 units. **(a)** Position-variant spiking rates (black) were fitted with skewed von Mises equations (red). **20%** peak amplitude in the fitted curve was used to define the beginning and end of place fields (red circles). Two examples were shown with narrow and wide fields, respectively. **(b)** Rate map of all 129 cells included in Fig.4 (under the control condition). Spiking rate of each cell was normalized by its peak. Cells were sorted by the center of mass of their place fields. For each cell, white and magenta ticks marked the beginning and end of its place field, determined by the method described in a. **(c)** Distribution of place field width. The median field width is 36.4cm.

